# Human PrP E219K as a new and promising substrate for RT-QuIC amplification of human prion strains: a first step towards strain discrimination

**DOI:** 10.1101/2024.07.31.605973

**Authors:** A. Marin-Moreno, F. Reine, F. Jaffrézic, L. Herzog, H. Rezaei, I. Quadrio, S. Haïk, V. Béringue, D. Martin

**Affiliations:** Université Paris-Saclay, Institut National de Recherche pour l’Agriculture, l’Alimentation et l’Environnement, Université Versailles-Saint Quentin, Unité de Virologie et d’immunologie Moléculaires, UMR 892, Jouy-en-Josas, France; Laboratorio Central de Veterinaria, Ministerio de Agricultura, Pesca y Alimentación, Algete (Madrid), Spain; Université Paris-Saclay, Institut National de Recherche pour l’Agriculture, l’Alimentation et l’Environnement, AgroParisTech, Unité de Génétique Animale et Biologie Intégrative, Jouy-en-Josas, France; Hôpital neurologique, Lyon, France, Biochemistry and Molecular Biology Department, Neurodegenerative Pathologies, LBMMS, 59 boulevard Pinel, 69500 Bron, Hospices Civils de Lyon, Lyon, France; BIORAN Team, Lyon Neurosciences Research Center, 59 boulevard Pinel, 69500 Bron, CNRS UMR 5292, INSERM U1028, Lyon, France; Sorbonne Université, Institut du Cerveau - Paris Brain Institute - ICM, Inserm, CNRS, Paris, France

**Keywords:** Prion, Creutzfeldt-Jakob disease, RT-QuIC, prion amplification, strain discrimination

## Abstract

Prion diseases are fatal neurodegenerative diseases that affect mammals through the transconformation of a host protein, the prion protein (PrP), into a toxic and pathogenic conformer termed PrP^Sc^. Until now, the diagnosis is only confirmed with a post-mortem histology study of the central nervous system. Among the methods to detect the etiological agent, *in vitro* amplification techniques have emerged as very sensitive, highly specific and rapid tools, even though some prion strains remain refractory or difficult to amplify. Here we report the use of a new recombinant substrate for Real-Time Quaking Induced Conversion (RT-QuIC), a natural polymorphism of human prion protein with a lysine at position 219 instead of a glutamic acid, PrP E219K. This substrate amplifies the six sporadic human strains responsible for Creutzfeldt-Jakob Disease (CJD) and the strain responsible for its variant form in a few hours and over a large dilution range of the seeds. Moreover, based on the lag time of the amplification reactions, the PrP E219K substrate allows to discriminate between sporadic and variant CJD strains, a first step towards an ante-mortem typing of the prion strain affecting a patient.

## Introduction

Transmissible Spongiform Encephalopathies (TSE), commonly called prion diseases, are fatal neurogenerative diseases. TSE can affect mammals including ruminants (sheep, cow, cervid) and Humans[15, 32, 34] and can be experimentally transmitted to rodents (mouse, hamster, bankvole)[36]. In Humans, Creutzfeldt-Jakob disease (CJD) is the most prevalent prion disease. Sporadic CJD (sCJD) is the most common CJD form, with approximately 1.5 cases per million and year, representing 80% of TSE in Humans[31]. Other human TSE can be either genetic like Fatal Familial Insomnia (FFI) and Gerstmann-Sträussler-Scheinker Syndrome (GSS) or acquired like variant CJD (vCJD) which is related to the consumption of food contaminated by prions responsible for Bovine Spongiform Encephalopathy[14]. Prion diseases are due to the transconformation of the host cellular prion protein (PrP^C^) into infectious conformers termed PrP^Sc^ [47–49]. PrP^Sc^ aggregates into oligomers and amyloids, that mainly accumulate in the central nervous system. PrP^Sc^ assemblies are more resistant to protease degradation than PrP^C^ and the resistant core of the protein (PrP^res^) is a biochemical signature of the disease.

Different prion strains are identified in defined hosts, each exhibiting unique phenotypes, including disease duration, clinico-pathological presentation and PrP^res^ electrophoretic signature[5]. These traits are encoded within structurally distinct PrP^Sc^ conformers. Human sCJD comprises a broad spectrum of clinicopathological variants, due to the combined existence of multiple strains and host polymorphism at codon 129 in the PrP encoding gene *PRNP*, that encodes either methionine (M) or valine (V). Depending on the molecular weight of its unglycosylated isoform, the PrP^res^ present in sCJD-infected individuals is classified as type 1 (21-kDa fragment) or type 2 (19-kDa fragment)[6, 44]. According to molecular PrP^res^ type genotype at codon 129 and histopathological phenotype, sCJD is thus classified at least into 7 molecular subtypes: MM1, MV1, VV1, MM2 (thalamic and cortical), MV2 and VV2. Experimental transmissions of cases representative of these subtypes to various lines of transgenic mice expressing human PrP (*humanized* mice) allow demonstrating that different strains were (at least partially) associated to these subtypes[40]. According to Bishop *et al*. categorization, using codon 129 genotype and PrP^Sc^ electrophoretic signature in the donor brain, M1 and V2 strains are the most prevalent, responsible for MM1 and MV1 cases, and VV2 and MV2 cases, respectively. V1 strain is responsible for VV1 cases, M2c and M2t for MM2 cortical and thalamic forms, respectively. Depending of the humanized line used, VV2 and MV2 could lead to different biological phenotype, suggesting different strains[58]. Co-propagation of M1 and V2 in the brains of sCJD-infected individuals is also observed in a large number of MM1/MV1 and MV2/VV2 cases[10].

CJD diagnosis is mainly based on evidence for rapidly deteriorating neurogenerative symptoms, dementia and Magnetic Resonance Imaging of the patient’s brain[26]; however, definitive confirmation is only performed with post-mortem histological and biochemical studies[8]. However, situation may change in the near future thanks to *in vitro* amplification techniques that are very efficient in amplifying minute amounts of PrP^Sc^ assemblies. Protein Misfolding Cyclic Amplification (PMCA) uses as a substrate PrP^C^ from healthy brains, mostly from transgenic mice expressing different types of mammalian PrP^C^ [52]. Variant CJD is well amplified using PMCA compared to MM1 which is rather refractory to PMCA[9, 20]. The second *in vitro* amplification technique is Real-Time Quaking Induced Conversion (RT-QuIC) which uses recombinant PrP expressed and purified from *E. Coli*[1–3]. All sCJD strains are well amplified using this method, however, vCJD is often reported as more difficult to amplify compared to MM1 sCJD[42, 45]. Moreover, since RT-QuIC amplified products are barely infectious, RT-QuIC is believed to unfaithfully amplify prion strains, losing the PrP^res^ signature of the prion seed[53], and preventing prion strain discrimination[1, 21].

RT-QuIC has several advantages over PMCA. First it allows the detection in real time, using a fluorophore (Thioflavin T), whose fluorescence parameters change upon its binding to amyloid structures. Usually within 48 hours of amplification reaction, prion detection can be achieved using brain homogenates as seeds. RT-QuIC is hence faster than PMCA and does not require several manipulation steps (several rounds of reaction, Proteinase K digestion to remove the excess of PrP^C^, and western-blot detection of PrP^res^). RT-QuIC is thus more convenient for high-throughput experiments dedicated to diagnosis or anti-prion drug screening for example[11]. Due to the low if any infectivity of RT-QuIC mixtures, this technique has lower biohazard issues compared to PMCA. It is also highly versatile in terms of source of seeds (cerebrospinal fluid (CSF), blood, nasal brushes, saliva, urine, feces, skin, muscles, eye drops…)[3, 7, 13, 17, 25, 29, 33, 35, 41] and RT-QuIC using CSF was incorporated in the diagnostic criteria for sCJD of several surveillance centres[26, 27, 30, 59]. Yet, it does not differentiate between human strains. As mentioned above, strains determine the disease evolution. Also, due to the different biochemical properties of prion strains, it is known that the efficacy of anti-prion therapy towards human strains may be strain-specific[24]. In addition, since prion decontamination is a delicate harsh process and considering that experimental anti-prion therapies are more effective when administered early in the course of the disease, there is an urgent need to rapidly detect prions in patients to manage secondary transmission risks, especially after invasive surgeries, and to allow fast inclusion of patients and stratification in future clinical trials. Early detection (at the initial clinical stage of the disease or before onset of the disease in some at-risk populations) and strain discrimination are thus important issues to be addressed. The finding of a RT-QuIC substrate that meets both requirements: fast and robust detection of all human prion strains at the same time, including those reported as difficult to amplify and, when possible, strain discrimination, is of particular need.

Here we report the use of the E219K polymorphism of human PrP M129 allele (PrP E219K), a dominant-negative polymorphism reported in the asian population to protect against sCJD[55], as a relevant substrate to achieve this goal. Using this protein, expressed and purified from *E. Coli*, we were able to amplify all sCJD and vCJD strains, either after passage in human PrP M129 transgenic mice (tg650) or directly from patient’s brains, in less than 20 hours and with a dilution range of terminally ill tg650 brain homogenates (BH) and most patient brain homogenates usually spanning between 5 and 8 logs. Based on the lag time of the amplification reactions, PrP E219K allows easy discrimination between sporadic and variant CJD. Moreover, sCJD MM2-c strain after passage and adaptation to tg650 mice could also be discriminated from the other sCJD strains. Using this substrate, we were able to differentiate MM1 and MV1 strains after passage and adaptation to tg650 mice, despite their similar phenotypes. These results support the view that PrP E219K is a versatile substrate of choice to achieve fast and robust detection and amplification of human prions.

## Materials and Methods

### Brain homogenates (BHs) from infected tg mice and Western-blot

PrP^res^ was extracted from 20% BHs with the Bio-Rad TeSeE detection kit, as previously described[4, 12]. Briefly, 200 μL aliquots were digested with proteinase K (200 μg/mL final concentration in buffer A) for 10 min at 37°C before precipitation with buffer B and centrifugation at 28,000 × *g* for 5 min. Pellets were resuspended in Laemmli sample buffer, denatured, run on 12% Bis-Tris Criterion gels (Bio-Rad, Marne la Vallée, France), electrotransferred onto nitrocellulose membranes, and probed with 0.1 μg/mL biotinylated anti-PrP monoclonal antibody Sha31 antibody (human PrP epitope at residues 145 to 152)[16], followed by streptavidin conjugated to horseradish peroxidase (HRP). Immunoreactivity was visualized by chemiluminescence (Pierce ECL, Thermo Scientific, Montigny le Bretonneux, France). The size and relative amounts of PrP^res^ glycoforms were determined using Image Lab software after the acquisition of chemiluminescent signals with the Chemidoc digital imager (Bio-Rad, Marne la Vallée, France).

### Real Time-Quaking Induced Conversion (RT-QuIC)

RT-QuIC amplifications were performed as previously described[37, 39]. In brief, 1 µL of 10% human BHs or 20% mouse BHs underwent serial 10-fold dilution in 20 mM sodium phosphate buffer pH 7.4, 130 mM NaCl, 0.1% SDS and 1X N2 supplement (Thermo Fisher, France). Then, 2 µL of each dilution was loaded in individual wells of a black 96-well optical bottom plate containing 98 µL of 20 mM sodium phosphate buffer pH 7.4, 300 mM NaCl, 10 µM thioflavin-T, 1 mM EDTA and 100 µg/mL of purified recombinant E219K human PrP polymorphism (harbouring the M_129_ allele)[50]. The plate was sealed with Nunc Amplification Tape (Nalgene Nunc International, France), placed in a Xenius XM spectrofluorometer (Safas, Monaco), and incubated for 24–60 h at 46°C ± 2°C. Until the end of the measurements, cycles of 1-min orbital shaking (600 rpm) and 1-min rest were applied, with fluorescence recorded every 30 min. The experiments were performed in quintuplicate. Each curve was fitted with the following equation,

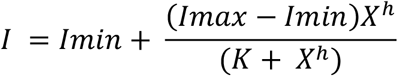

using MATLAB (R2022b, MathWorks) where I is the fluorescence intensity and X is the time. The parameters calculated from the fit included the increase in fluorescence intensity, the slope at the inflection point and the lag time (estimated by extending the tangent at the inflection point to the initial baseline Ymin)[37, 39]. The seeding activity titer (SD_50_; seeding dose giving thioflavin-T positivity in 50% of the replicates) was estimated using the Spearman– Kärber method[38]. When <100% of the RT-QuIC reactions seeded with the first dilution scored positive or when no dilution scored 100% positive, a trimmed variant of the Spearman– Kärber method was applied[23]. The titer was expressed as SD_50_ per mL of 10% (w/v) BH. Statistical analyses on RT-QuIC significance were done using the nlme[46] and lmPerm[56] packages in R. For strain or isolate comparison, a linear mixed-effects model and a permutation test for linear models were used to compare the area under the lag time curves. The models included as a fixed effect the prion strain. The Tukey method has been used for P value adjustment of multiple comparison.

## Results

### PrP E219K RT-QuIC amplifies tg650-adaptated sporadic CJD prions

We first assayed PrP E219K as a substrate in RT-QuIC amplification experiments using brain homogenates (BHs) from terminally ill humanized tg650 mice (PrP M129 allele) which were iteratively inoculated with MM1, MV1, MM2-c, MV2, VV2, VV1 sCJD cases (4 to 5 passages)[12, 22, 28]. For clarity, tg650 adapted strains are named tg650-MM1, tg650-MV1, tg650-VV1, tg650-MM2c, tg650-MV2 and tg650-VV2. In these mice, MM1 and MV1 cases produced similar phenotype, suggesting infection by the same M1 strain. The other cases lead to different and unique phenotypes, suggesting at least isolation of 5 strains in these mice. BH of healthy tg650 mice was used as a negative control. We performed 10-fold serial dilutions (spanning 8 logs) of the initial 20% BH in the seed dilution buffer (PBS-SDS-N2). Two uL of these dilutions were then used to initiate RT-QuIC reactions. As previously reported in[37, 39], the fitted fluorescence intensity mean of the positive reaction at each dilution for each strain as a function of time is presented in supplementary Figure 1. From these curves, the lag time was calculated for each positive reaction. Moreover, from the number of positive reactions at each dilution and using the Spearman-Kärber equation, we estimated a SD50 and the concentration of amplifiable particles in the samples for each strain (Table 1). Figure 1A shows the lag time as a function of particle concentration.

**Table 1:** quantifications of CJD strains after tg650 passage and adaptation. Dilution expressed from a 100% brain homogenate, from which 2 μL were used to seed the amplification reaction. SD50 is estimated as explained in materials and methods. ID50 has been determined after inoculation and adaptation to tg650 mice (nd: not determined). The quantification of PrP^Sc^ has been estimated by western-blot as explained in materials and methods.

| Strain | Tg650-MM1 | Tg650-MV1 | Tg650-VV1 | Tg650-MM2 | Tg650-MV2 | Tg650-VV2 | Tg650-vCJD |
| --- | --- | --- | --- | --- | --- | --- | --- |
| Dilution range | > 7 log | 5 log | 7 log | 6 log | 5 log | 6 log | 6 log |
| 1<sup>st</sup> positive dilution | 2.10<sup>-3</sup> | 2.10<sup>-3</sup> | 2.10<sup>-2</sup> | 2.10<sup>-3</sup> | 2.10<sup>-3</sup> | 2.10<sup>-3</sup> | 2.10<sup>-3</sup> |
| Last positive dilution | > 2.10<sup>-9</sup> | 2.10<sup>-7</sup> | 2.10<sup>-8</sup> | 2.10<sup>-8</sup> | 2.10<sup>-7</sup> | 2.10<sup>-8</sup> | 2.10<sup>-8</sup> |
| SD50 / mg of brain | 7,91.10<sup>8</sup> | 4,45.10<sup>7</sup> | 4,45.10<sup>7</sup> | 2,50.10<sup>7</sup> | 1,47.10<sup>7</sup> | 7,91.10<sup>7</sup> | 7,91.10<sup>7</sup> |
| ng PrP<sup>Sc</sup> / mg of brain | 18,2 | 7,7 | 32,9 | 4,5 | 45,6 | 36,3 | 860,4 |
| ID50 / mg of brain | 5,01.10<sup>6</sup><br>[39] | nd | nd | 1,26.10<sup>5</sup><br>[12] | 2,29.10<sup>5</sup> | nd | 6,31.10<sup>5</sup><br>[22] |

**Figure 1.**
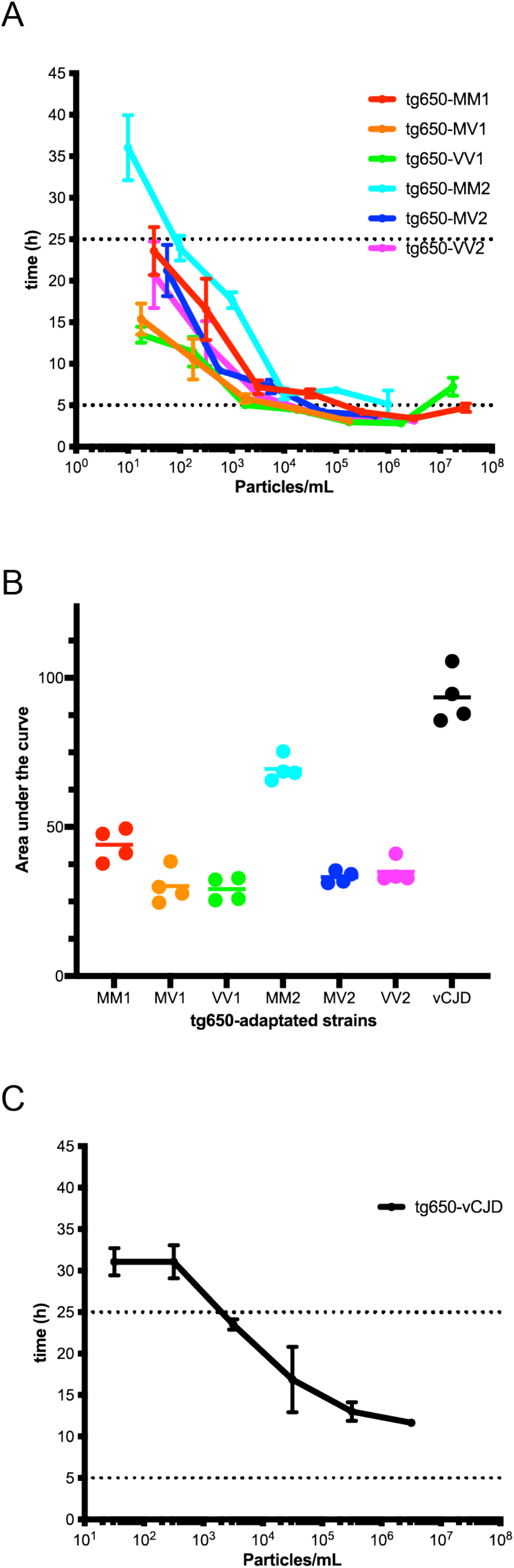
Analyses of infected transgenic mouse brain homogenates. A) lag time of amplification reactions seeded with tg650-sCJD BHs. B) Area under the curve as a function of the strain. C) lag time of amplification reactions seeded with tg650-vCJD BH.

As shown on supplementary Figure 2, no auto-polymerization of PrP E219K was observed over the duration of the amplification reaction (50 hours). Figure 1A and Supplementary Fig 1 show that the 6 tg650-sCJD prions could be amplified using PrP E219K as a substrate over a large dilution range: from at least 5 log dilution for tg650-MV1 and tg650-MV2 to more than 7 log dilution for tg650-MM1 (Table 1). Except for tg650-VV1, we observed an inhibition of the amplification reactions at the first dilution corresponding to a 500-fold dilution of BHs, as commonly reported in the literature[43]. From the lag time as a function of particle concentration, we observed that the lag time increased with the dilution, except for concentrations above 10^7^ particles per mL due to the inhibition mentioned above. Except for the limit dilution with tg650-MM2-c prions, all reactions started before 25 hours; the lag time was generally around 5 hours for all strains covering at least a 3 or 4 log dilution range. To compare the tg650-strains, the area under the curve (AUC) of the lag time as a function of the particle concentration for each strain and on the same amplification range was calculated (Figure 1B). Statistical analyses on pairwise comparisons were performed using two models, a parametric linear mixed-effects model and a non-parametric permutation test for linear models. Both analyses (Table 2) show that tg650-MM2-c sCJD prions strain can be discriminated from the other tg650-sCJD prions strains. Moreover, our results show that even if tg650-MM1 and tg650-MV1 share a similar phenotype, they do not have the same seeding activity (Table 2) and thus can be differentiated based on RT-QuIC amplification reactions. Collectively, these data suggest that PrP E219K substrate can amplify several human prion strains and with a certain degree of discrimination based on the lag time of the amplification reactions.

**Table 2:** P values of statistical analyses concerning strain comparison (after adaptation to tg650 mice) using both a parametric linear model and a non-parametric linear model with permutations.

|  | Tg650-MV1 | Tg650-VV1 | Tg650-MM2 | Tg650-MV2 | Tg650-VV2 | Tg650-vCJD |
| --- | --- | --- | --- | --- | --- | --- |
| Tg650-MM1 | 0.0195 | 0.0106 | < 0.0001 | 0.1022 | 0.2444 | < 0.0001 |
| Tg650-MV1 | / | 1.0000 | < 0.0001 | 0.9826 | 0.8507 | < 0.0001 |
| Tg650-VV1 |  | / | < 0.0001 | 0.9290 | 0.7064 | < 0.0001 |
| Tg650-MM2 |  |  | / | < 0.0001 | < 0.0001 | < 0.0001 |
| Tg650-MV2 |  |  |  | / | 0.9988 | < 0.0001 |
| Tg650-VV2 |  |  |  |  | / | < 0.0001 |

**Figure 2.**
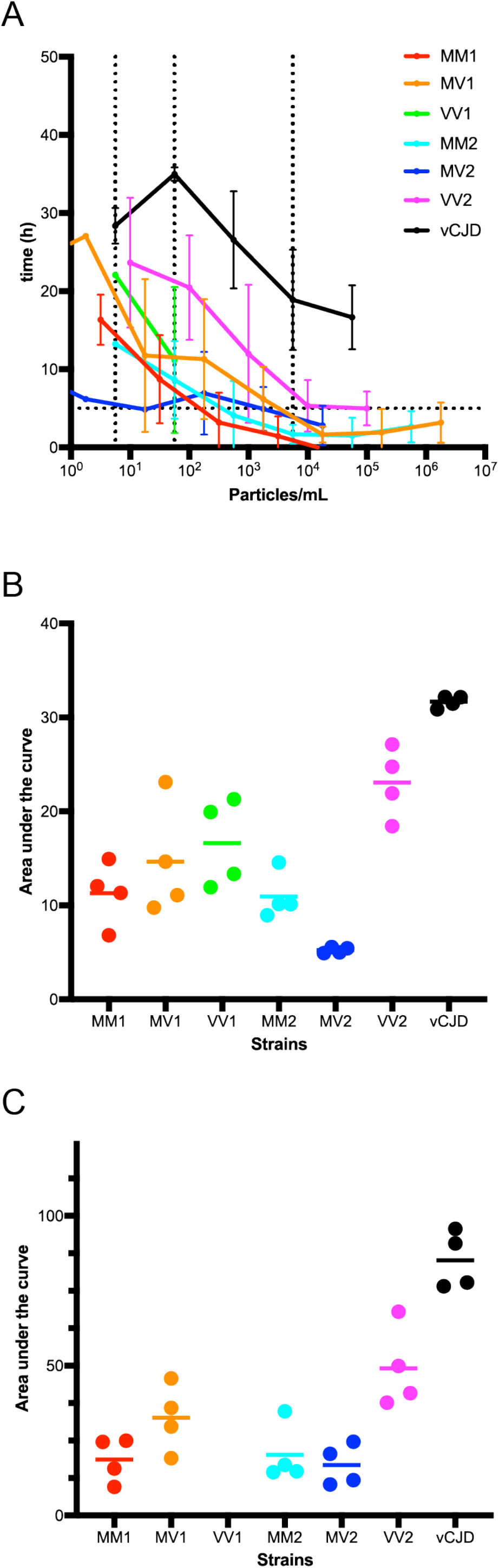
Analyses of human brain homogenates. A) lag time of sCJD and vCJD seeded reactions with patient brain homogenates. B) Area under the curve as a function of the strain over 2-log dilution. C) Area under the curve as a function of the strain over 4-log dilution.

### PrP E219K RT-QuIC discriminates tg650-variant CJD prion from tg650-sCJD prions

We next examined if PrP E219K could amplify a tg650-adapted variant CJD (tg650-vCJD), derived from zoonotic BSE prions. We used 10-fold serial dilutions (spanning 8 logs) of the initial 20% BH of terminally ill tg650 mice infected with the biologically cloned vCJD prion strain[22]. The fitted fluorescence intensity mean of the positive reactions at each dilution for each strain as a function of time is presented in supplementary Figure 1C. Supplementary Figure 3 and Figure 1C show that tg650-vCJD prions could be readily amplified using PrP E219K as a substrate. The dilution range spans over 6 logs for tg650-vCJD strain (Table 1). The amplification reactions started between 11 and 31 hours, and the lag time was always longer for tg650-vCJD than for tg650-sCJD at a given seed concentration. The same statistical analyses of the AUC of the lag time as a function of the particle concentration was performed as above (Figure 1B, Table 2). Based on these analyses and on this parameter of the kinetics, PrP E219K allowed to discriminate between tg650-sCJD and vCJD prions.

### PrP E219K RT-QuIC amplifies sCJD and vCJD prions directly from patient brain homogenates

We next wondered if PrP E219K could be used to amplify human prions from BHs of s-and v-CJD patients. To investigate this, we used the human BHs that were used as inocula to infect tg650 mice. We also had access to a human healthy BH that we used as negative control. We performed 10-fold serial dilution (spanning 9 logs) of the initial 10% BHs to seed the RT-QuIC reactions. The fitted fluorescence intensity mean of the positive reaction at each dilution for each sample as a function of time is presented in supplementary Figure 4. As shown on supplementary Figure 4, no auto-polymerization of PrP E219K was observed over the duration of the amplification reactions (40 hours) when the reaction was seeded by the dilutions of the healthy BH. All CJD patient BHs gave positive amplification reactions using PrP E219K as a substrate (supplementary Figure 4 and Table 3). Except for the patient typed as VV1, all sCJD patient BHs gave positive reactions over a large dilution range (at least 5 logs). For the vCJD case, the dilution range spanned over 5 logs. From these curves, the lag time was calculated for each positive reaction. Moreover, from the number of positive reactions at each dilution and using the Spearman-Kärber equation, we estimated a SD50 and the concentration of amplifiable particles in the samples for each strain. Figure 2A shows the lag time as a function of particle concentration. As previously observed with the tg650-adaptated prions, human vCJD prion had a longer lag time than sCJD prions, around 25 hours for vCJD compared to usually less than 10 hours for all sCJD prions (Figure 2A). To further compare these prions, the AUC of the lag time as a function of the particle concentration and on the same amplification range was calculated (Figure 2 B and C). Two analyses were performed, one covering the short dilution range of VV1 prions (Figure 2B) and a second one covering a larger dilution range but excluding VV1 (Figure 2C). Statistical analyses on pairwise comparisons were performed using two models, a parametric linear mixed-effects model and a non-parametric permutation test for linear models. Both analyses (Table 4) show that based on the lag time parameter, vCJD prion can be discriminated from the other sCJD prions. These results confirm that PrP E219K is a relevant RT-QuIC substrate to amplify both sporadic and variant CJD prions and that the lag time of human prion amplification reaction using PrP E219K as a substrate is a relevant parameter to discriminate between sCJD and vCJD prions isolates.

**Table 3:** quantifications of CJD strains from human patients. Dilution expressed from a 100% HB, from which 2 μL were used to seed the amplification reaction. SD50 is estimated as explained in materials and methods.

| Strain | MM1 | MV1 | VV1 | MM2 | MV2 | VV2 | vCJD |
| --- | --- | --- | --- | --- | --- | --- | --- |
| Dilution range | 6 log | 8 log | 2 log | 6 log | 6 log | 5 log | 5 log |
| 1<sup>st</sup> positive dilution | 10<sup>-3</sup> | 10<sup>-3</sup> | 10<sup>-3</sup> | 10<sup>-3</sup> | 10<sup>-3</sup> | 10<sup>-3</sup> | 10<sup>-3</sup> |
| Last positive dilution | 10<sup>-8</sup> | 10<sup>-10</sup> | 10<sup>-4</sup> | 10<sup>-8</sup> | 10<sup>-8</sup> | 10<sup>-7</sup> | 10<sup>-7</sup> |
| SD50 / mg of brain | 1,58.10<sup>10</sup> | 8,89.10<sup>10</sup> | 2,81.10<sup>6</sup> | 2,81.10<sup>10</sup> | 8,89.10<sup>8</sup> | 5,00.10<sup>9</sup> | 2,81.10<sup>9</sup> |

**Table 4:** P values of statistical analyses concerning strain comparison (human BHs). Analyses were performed using both a parametric linear model and a non-parametric linear model with permutations. The first lines correspond to analyses of AUC over a 2-log dilution, the second ones to analyses of AUC over a 4-log dilution.

|  | MV1 | VV1 | MM2 | MV2 | VV2 | vCJD |
| --- | --- | --- | --- | --- | --- | --- |
| MM1 | 0.8306 | 0.3835 | 1.0000 | 0.2525 | 0.0023 | < 0.0001 |
|  | 0.3925 | / | 0.9999 | 0.9998 | 0.0048 | < 0.0001 |
| MV1 | / | 0.9846 | 0.7628 | 0.0185 | 0.0437 | < 0.0001 |
|  | / | / | 0.5139 | 0.2696 | 0.2330 | < 0.0001 |
| VV1 |  | / | 0.3167 | 0.0032 | 0.1963 | 0.0001 |
|  |  | / | / | / | / | / |
| MM2 |  |  | / | 0.3106 | 0.0017 | < 0.0001 |
|  |  |  | / | 0.9964 | 0.0077 | < 0.0001 |
| MV2 |  |  |  | / | < 0.0001 | < 0.0001 |
|  |  |  |  | / | 0.0028 | < 0.0001 |
| VV2 |  |  |  |  | / | 0.0368 |
|  |  |  |  |  | / | 0.0009 |

## Discussion

RT-QuIc is currently emerging among the most relevant assay for the pre-mortem diagnosis of sporadic CJD, with multiple applications for disease surveillance and differential diagnosis from other rapidly progressive dementia, limiting iatrogenic risks, identification of tissues at risk and therapeutic trials at the prodromal phases. However, the heterogenous phenotypic spectrum of sCJD due to host genetic factors and the existence of several prion strains is a potential limitation of the diagnostic performance and clinical utility of RT-QuIC, particularly when rarer subtypes of sCJD are considered. In addition, the assay offers little value in the differential diagnosis of sporadic CJD and variant CJD, the prions responsible for vCJD being amplified with limited sensitivity. While the number of clinical cases of vCJD epidemic remain limited, there are great uncertainties regarding the number of asymptomatic carriers of the disease[18], necessitating the development of diagnostic markers of the infection, including in peripheral tissues.

The four major recombinant PrPs used as RT-QuIC substrates in the literature are hamster PrP, human PrP, a chimeric hamster-human PrP and a truncated hamster PrP. Here, we report recombinant human PrP E219K, produced in *E. Coli*, as a promising RT-QuIC substrate to rapidly amplify and detect human sCJD subtypes and vCJD prions.

RT-QuIC allows the amplification of minute amounts of prions, however due to stochastic and spontaneous auto-polymerization of the substrate, discrimination between positive seeded reactions and negative ones related to auto-polymerization is of particular importance. The reactions seeded with dilutions of sane BHs, either from transgenic mice or of human origin, showed that PrP E219K did not auto-polymerise, during the time of our amplification experiments (up to 50 hours), the number of false positive reactions was thus null. The monitoring of fluorescence increase was thus related to a positive amplification reaction of prions. The analysis of the positive and negative samples was hence easy and fast compared to amplification using other recombinant proteins[19, 54, 57].

Our experiments highlight that PrP E219K could amplify all sCJD subtypes and vCJD prion, be there from tg650-adaptated or human origins directly. Except VV1, all tested human prions could be amplified over a large dilution range, at least 5 logs, and usually between 6 and 7 logs. For VV1, the dilution range of seeds from tg650 mouse HBs was 7 logs compared to 2 logs for seeds from the patient’s BH. Further investigations, *i*.*e*. more VV1 cases and pieces of information concerning the VV1 sample (like sampled brain area, treatment before homogenisation, conservation, or patient stage), would be necessary to address this apparent discrepancy, however, a marked decrease of RT-QuIC sensitivity has already been reported in the literature for MM2 and VV1 sCJD strains[51] and may explain our result. Testing a larger cohort of samples of various strains would be interesting to investigate whether PrP E219K would improve the diagnosis of these less prevalent and detectable strains. According to our results, PrP E219K is not sensitive to the brain matrix, since we were able to perform prion amplification directly from both tg650 and human BHs.

In this study, we performed limit dilution measurements to estimate the amount of each prion that we could detect and amplify using PrP E219K as substrate and direct dilutions of brain homogenates as PrP^Sc^ source. We estimated the amount of PrP^res^ per mg of brain by western-blot (Table 1). The amount of seeds that we could detect in the 2 uL of sample to seed the reaction was around 1 fg for VV1, VV2 and MV2, around 0.2 fg for MV1 and MM2, below 0.05 fg for MM1 and around 30 fg for vCJD. The values reported in the literature are around 1 fp for sCJD MM1, MM2 and MV2[3, 43, 45] and around 100 fg for vCJD[45] using other substrates (*i*.*e*. hamster PrP, human PrP or a chimeric hamster-human PrP). Our detection limits are thus in agreement with those previously reported, and even 5 times lower for vCJD, MV1, MM2-c and more than 20 times lower for MM1. We emphasize that we obtained these detection limits directly using brain homogenates without the need to capture and concentrate the seeds. In order to lower the detection limit, steps to concentrates the seeds, like immunoprecipitation, have been described[42]. However, increasing the number of steps increase the time of manipulation and error sources. Moreover, it is sometimes necessary to perform a second amplification round, by adding new substrate, again another error source, and increasing the reaction time could result in auto-polymerization and making more difficult the analysis of the results.

The lag time calculated from the kinetics stresses that PrP E219K allows to rapidly detect human prions. The majority of the sCJD seeded reactions started in a few hours (around 5 hours for concentrations higher than 10^3^ particles/mL). Our sCJD amplification reactions generally start 2 or 3 hours before what is reported in the literature using hamster or human PrP as substrate[3, 42, 45]. With human PrP, a lag time of 7 to 20 hours is usually observed. As far as vCJD seeded reactions are concerned, the reaction lag time was a bit longer (around 20 hours) to the ones of any sCJD amplification reactions. This observation is consistent to what is reported in the literature. However, when using PrP E219K as an amplification substrate, the lag time is still shorter than the lag time observed in the literature to amplify vCJD (above 30 hours with Hamster PrP for example). Shortening the lag time of the reaction has two advantages. First, the results are obtained more rapidly, saving time. Second, when the reaction time increase, auto-polymerization is more likely to occur. Longer lag time lowers the sensitivity of the technics and makes more hazardous the analysis of the data. More replicates are thus necessary to discriminate between true or false reaction. Moreover, based on the lag time of the amplification reactions, PrP E219K enables to clearly discriminate between sCJD and vCJD seeds. Of note, we also were able to discriminate tg650-MM2-c from the other tg650-sCJD strains, and we show that tg650-MM1 and tg650-MV1 which give similar phenotypes in tg650 mice have different seeding activities and are thus likely to be associated with different prion strains. RT-QuIC kinetic parameters and conditions appear as tools to discriminate between prion strains, as shown here or as shown to discriminated between tg650-adaptated MM1 and atypical scrapie[37]. Testing a lager cohort of samples and RT-QuIC conditions would be of interest to better achieve prion strain discrimination.

In conclusion, PrP E219K is a robust RT-QuIC substrate to rapidly detect and amplify all human CJD prion strains, and to easily discriminate between sCJD and vCJD according to the lag time parameter of the amplification reactions.

## List of abbreviations

AUC: Area Under the Curve
BH: Brain Homogenate
BSE: Bovin Spongiform Encephalopathy
CJD: Creutzfeldt-Jakob Disease
CSF: Cerebrospinal Fluid
E: glutamic acid
FFI: Fatal Familial Insomnia
GSS: Gerstmann-Sträussler-Scheinker Syndrome
K: lysine
M: methionine
N2: N2 supplement
PBS: Phosphate Buffer Saline
PrP: Prion Protein
PrP^C^: Cellular form of PrP
PrPres: protease-resistant core of PrP^Sc^
PrP^Sc^: Pathogenic conformer of PrP
RT-QuIC: Real-Time Quaking Induced Conversion
sCJD: sporadic Creutzfeldt-Jakob Disease
SDS: sodium dodecylsulfate
tg: Transgenic mouse
ThT: thioflavin-T
TSE: Transmissible Spongiform Encephalopathies
V: Valine
vCJD: variant form of Creutzfeldt-Jakob Disease

## Declarations

### Ethics approval and consent to participate

All animal experiments were approved by the INRAe Local Ethics Committee (Comethea; permit number 12/034). MM1-sCJD (UK NHBX0/0001), MV2-sCJD (UK NHBX0/0004) and vCJD (NHBY0/0003) samples were provided by the UK National Institute for Biological Standards and Control (NIBSC) CJD Resource Centre. MV1-sCJD Fr3 (A990055), VV2-sCJD Fr2 (A001002), VV1-sCJD Fr1 (367) and MM2-c-sCJD Fr1 (447) were provided by our collaborators (SH and IQ) within the frame of the French National Center of Reference for Unconventional Transmissible Agents and the French National Neuropathology Network for CJD, based on availability of autopsy-retained frozen brain material. For each case, an informed consent for genetic analysis of the PrP gene (*PRNP*) to obtain the genotype at codon 129 and to exclude the presence of pathogenic mutations was obtained from the patient’s relatives at the time of diagnosis. The patient’s relatives also gave written informed consent for autopsy and research using postmortem tissues in accordance with French regulations (L.1232-1 to L.1232-3, Code de la Santé Publique).

## Consent for publication

Not applicable

## Availability of data and materials

All the data generated and analyzed during this study are included in this published article and its supplementary information files.

## Authors’ contributions

A.M.M., V.B. and D.M. designed the research. A.M.M, F.R., F.J., L.H., H.R., S.H., I.Q., V.B. and D.M. performed the research and analyzed data. A.M.M, V.B. and D.M. wrote the paper. All authors approved the final version of the manuscript.

## Acknowledgement

We thank the staff of Animalerie Rongeurs (Infectiology of fish and rodent facility, Jouy-en-Josas, France) for animal care.

## Supplementary Figure Legends

Supplementary Figure 1: RT-QuIC amplification reactions of tg650-MM1, tg650-MM2c, tg650-MV1, tg650-MV2, tg650-VV1, tg650-VV2 as a function of brain homogenate dilution. The presented data are the mean (with standard deviation) of fitted positive amplification reactions.

Supplementary Figure 2: RT-QuIC reactions of tg650-healthy as a function of brain homogenate dilution. The presented data are the mean (with standard deviation) of fitted reactions.

Supplementary Figure 3: RT-QuIC amplification reactions of tg650-vCJD as a function of brain homogenate dilution. The presented data are the mean (with standard deviation) of fitted positive amplification reactions.

Supplementary Figure 4: RT-QuIC amplification reactions of MM1-sCJD, MM2c-sCJD, MV1-sCJD, MV2-sCJD, VV1-sCJD, VV2-sCJD, vCJD and healthy BH as a function of brain homogenate dilution. The presented data are the mean (with standard deviation) of fitted positive amplification reactions.

