## supplemental data for "Human PrP E219K as a new and promising substrate for RT-QuIC amplification of human prion strains: a first step towards strain discrimination"

**Supplementary Figure Legends**

Supplementary Figure 1: RT-QuIC amplification reactions of tg650-MM1, tg650-MM2c, tg650-MV1, tg650-MV2, tg650-VV1, tg650-VV2 as a function of brain homogenate dilution. The presented data are the mean (with standard deviation) of fitted positive amplification reactions.

Supplementary Figure 2: RT-QuIC reactions of tg650-healthy as a function of brain homogenate dilution. The presented data are the mean (with standard deviation) of fitted reactions.

Supplementary Figure 3: RT-QuIC amplification reactions of tg650-vCJD as a function of brain homogenate dilution. The presented data are the mean (with standard deviation) of fitted positive amplification reactions.

Supplementary Figure 4: RT-QuIC amplification reactions of MM1-sCJD, MM2c-sCJD, MV1-sCJD, MV2-sCJD, VV1-sCJD, VV2-sCJD, vCJD and healthy BH as a function of brain homogenate dilution. The presented data are the mean (with standard deviation) of fitted positive amplification reactions.

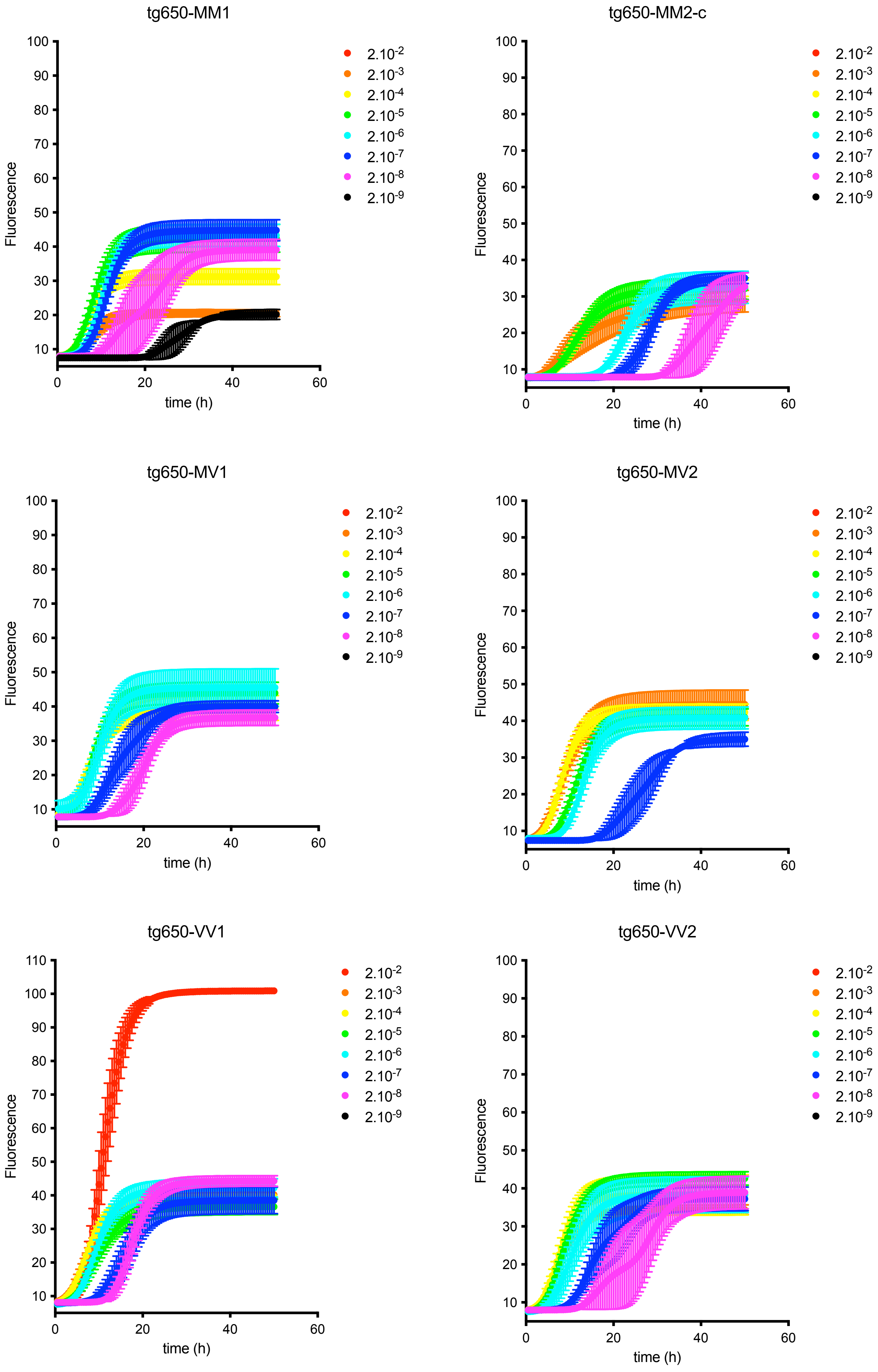
tg650-MM1
tg650-MM2-c
tg650-MV1
tg650-MV2
tg650-VV1
tg650-VV2
Fluorescence
time (h)
$2.10^{-2}$
$2.10^{-3}$
$2.10^{-4}$
$2.10^{-5}$
$2.10^{-6}$
$2.10^{-7}$
$2.10^{-8}$
$2.10^{-9}$

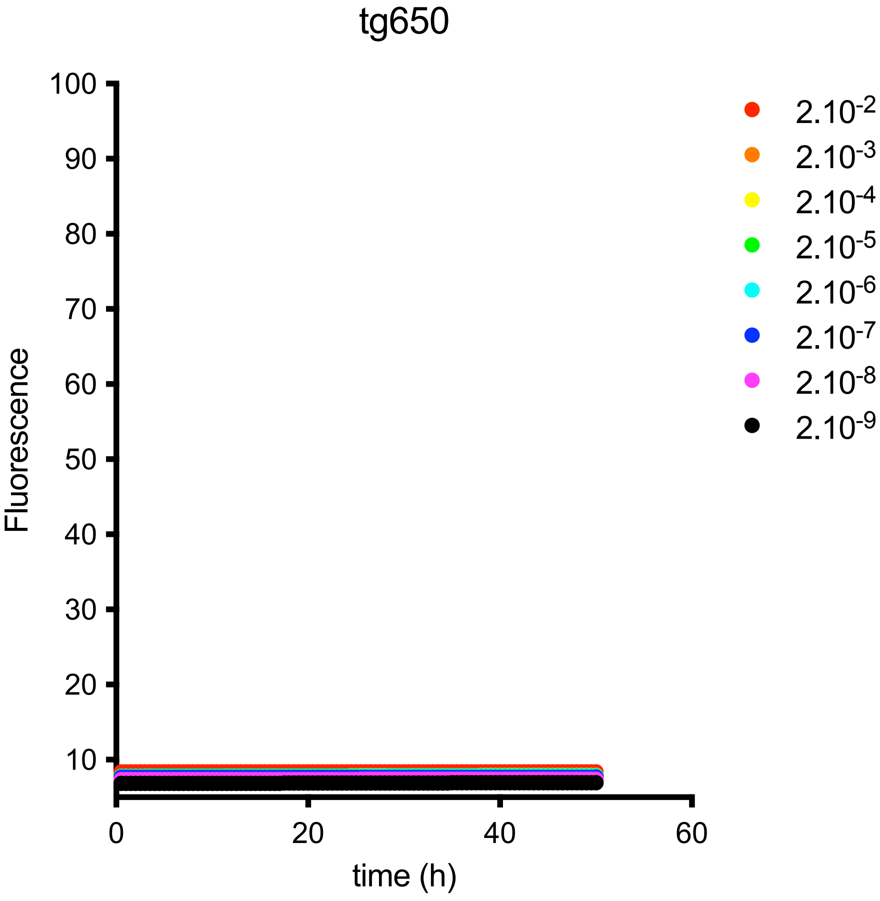

tg650
100
90
80
70
60
50
40
30
20
10
Fluorescence
0
20
40
60
time (h)
2.10-2
2.10-3
2.10-4
2.10-5
2.10-6
2.10-7
2.10-8
2.10-9

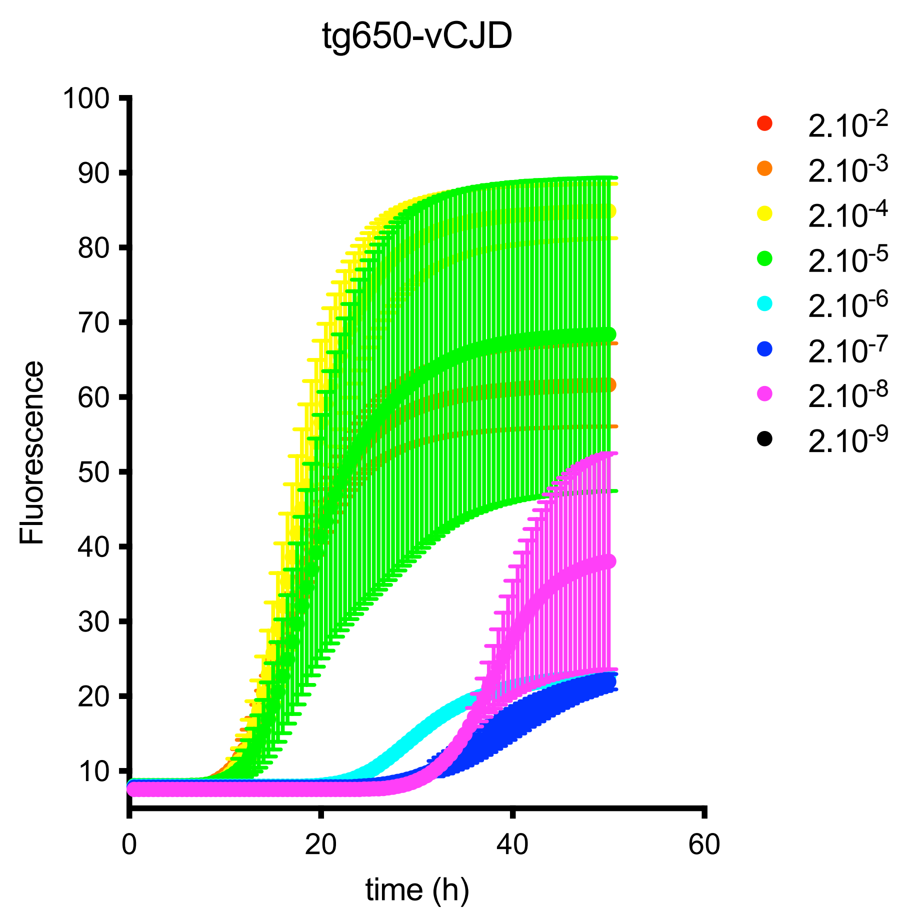

tg650-vCJD
Fluorescence
time (h)
0
20
40
60
10
20
30
40
50
60
70
80
90
100
2.10-2
2.10-3
2.10-4
2.10-5
2.10-6
2.10-7
2.10-8
2.10-9

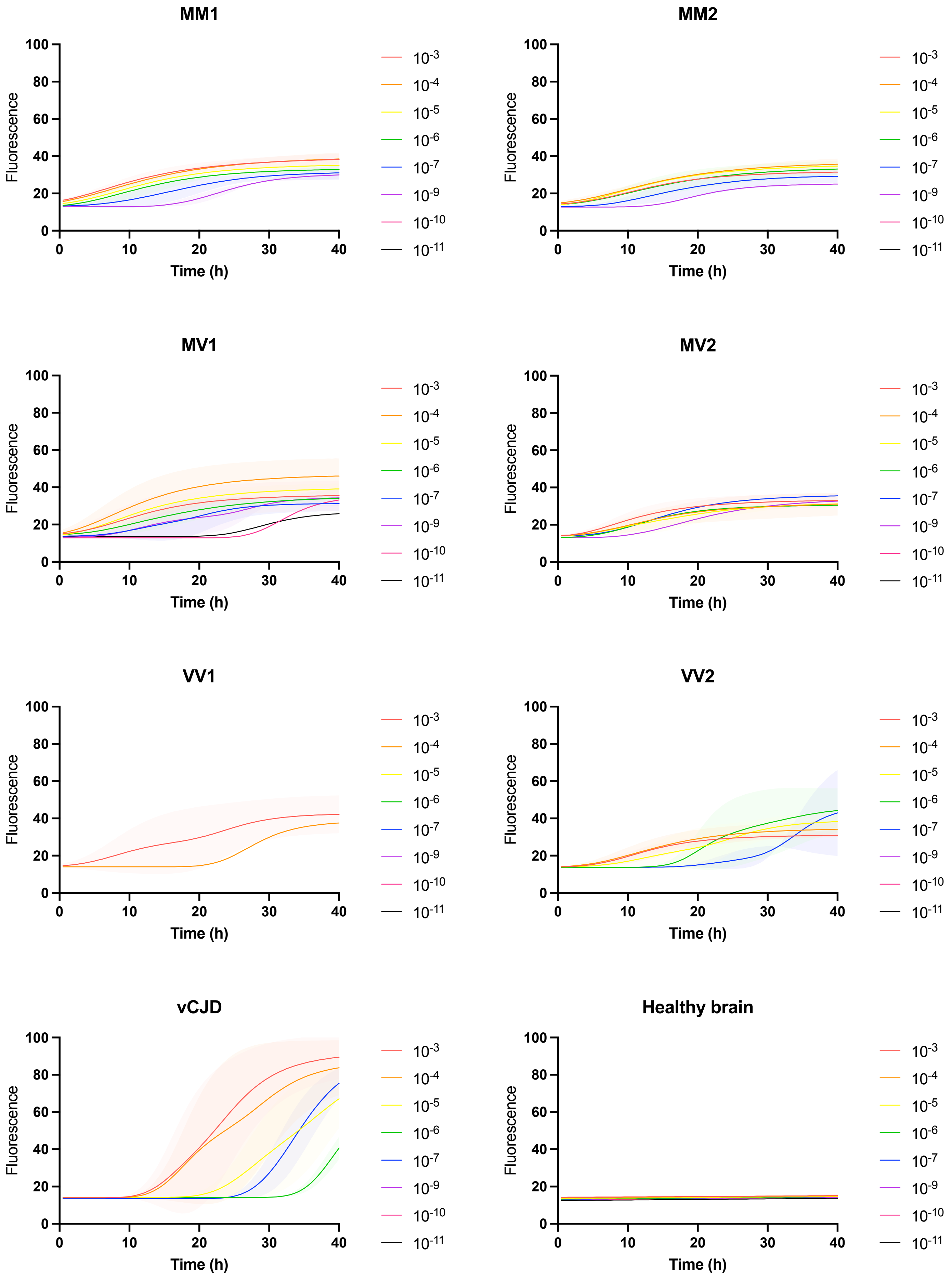
MM1
MM2
MV1
MV2
VV1
VV2
vCJD
Healthy brain
Fluorescence
Time (h)
0
20
40
60
80
100
10
30
$10^{-3}$
$10^{-4}$
$10^{-5}$
$10^{-6}$
$10^{-7}$
$10^{-9}$
$10^{-10}$
$10^{-11}$
